# Demographic tradeoffs predict tropical forest dynamics

**DOI:** 10.1101/808865

**Authors:** Nadja Rüger, Richard Condit, Daisy H. Dent, Saara J. DeWalt, Stephen P. Hubbell, Jeremy W. Lichstein, Omar R. Lopez, Christian Wirth, Caroline E. Farrior

## Abstract

Assessing vegetation feedbacks with the climate system and planning sustainable management in tropical forests requires efficient, yet accurate, predictions of the joint dynamics of hundreds of tree species. With increasing information on tropical tree life-histories, our predictive understanding is no longer limited by species data, but by the ability of existing models to make use of it. Using a demographic forest model, we show that the basal area and compositional changes during forest succession in a Neotropical forest can be accurately predicted by representing tropical tree diversity (hundreds of species) with only five functional groups spanning two essential tradeoffs – the growth–survival and stature–recruitment tradeoffs. This data-driven modeling framework substantially improves our ability to predict consequences of anthropogenic impacts on tropical forests.

## Main Text

Tropical forests are highly dynamic. Only about 50% of the world’s tropical forests are undisturbed old-growth forests (*1*). The remaining half comprises forests regenerating after previous land use, timber or fuelwood extraction, or natural disturbances. Even unmanaged old-growth forests are a dynamic mosaic of patches recovering from single- or multiple treefall gaps (*2*). Thus, understanding how forest structure and composition of the diverse tree flora change during recovery from disturbance is fundamental to predict carbon stocks and fluxes and vegetation feedbacks to the climate system (*3*), as well as to plan sustainable forest management (*4*). Despite the importance of regenerating tropical forests for the global carbon cycle and timber industry, our mechanistic understanding and ability to forecast compositional changes of these forests remains severely limited (*5*).

So far, forest succession has been viewed mostly through a one-dimensional lens distinguishing early-, mid-, and late-successional species (*5*, *6*). This classification corresponds to the well-known fast–slow life-history continuum or the growth–survival tradeoff (*7*). ‘Fast’ species grow quickly but survive poorly and dominate early successional stages, while ‘slow’ species grow slow but survive well and reach dominance in later successional stages. In addition, several studies suggest that tropical tree communities are also structured along a second major tradeoff axis that is orthogonal to the growth–survival tradeoff: the stature–recruitment tradeoff (*8, 9*). The stature–recruitment tradeoff distinguishes long-lived pioneers (LLPs) from short-lived breeders (SLBs). LLPs grow fast and live long and hence attain a large stature, but suffer low recruitment. SLBs grow and survive poorly and hence remain short-statured, but produce large numbers of offspring (*9*). However, we are lacking a systematic assessment of how important these tradeoffs are for predicting forest composition and carbon dynamics.

We parameterized a forest model based on height-structured competition for light (*10, 11*) with demographic tradeoffs derived from forest inventory data to uncover the essential number of demographic niches needed to accurately predict compositional changes during tropical forest succession. The model simulates the dynamics of a potentially large number of species based on a small set of demographic rates (growth, survival, recruitment) and accounts for height-structured competition for light by distinguishing up to four dynamic canopy layers (*12*). Canopy gaps are filled by the tallest trees from lower canopy layers, without regard for their horizontal position (*10*).

Our study site is the tropical moist forest at Barro Colorado Island (BCI), Panama, where recruitment, growth and survival of individual trees has been monitored in a 50-ha plot for over 30 years (*2, 12, 13*). To account for height-structured competition, we assigned all monitored individuals of 282 tree and shrub species to one of four canopy layers based on their size and the size of their neighbors (*12, 14*) and estimated model parameters (annual diameter growth and survival rates) for each species in each canopy layer (*9*). Additionally, we calculated species recruitment rates per unit of basal area. A dimension reduction of model parameters (weighted PCA, *15*) reveals the two demographic tradeoffs, i.e. the growth–survival tradeoff and the stature–recruitment tradeoff, which together explain 65% of demographic variation among the 282 species (Fig. 1).

**Fig. 1:**
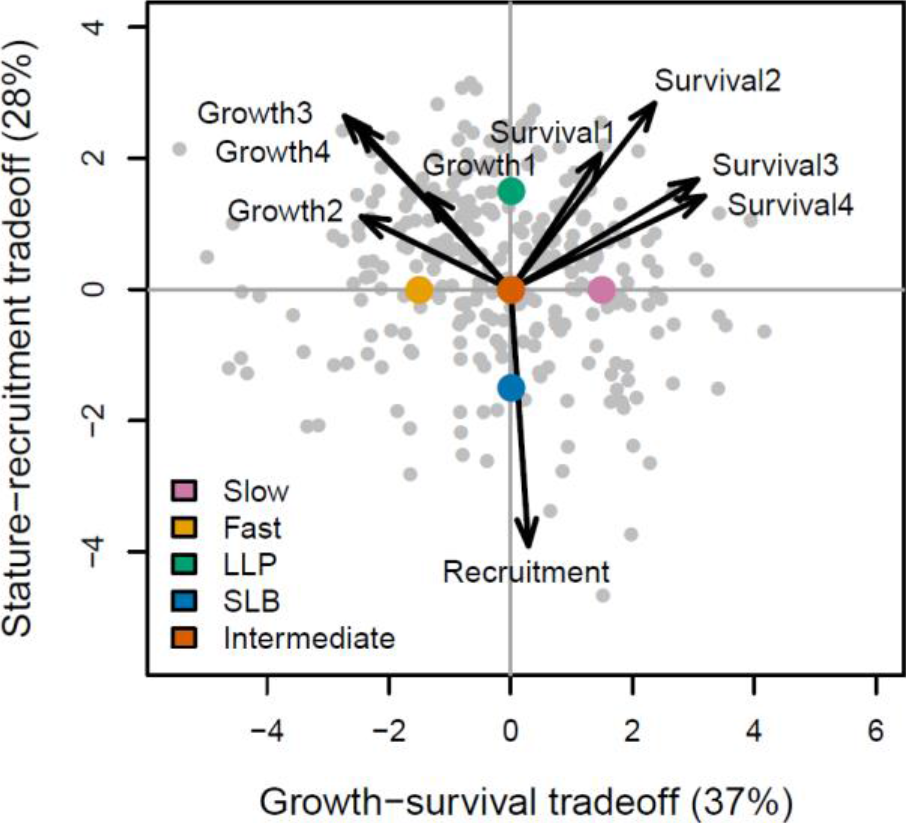
Demographic tradeoffs. Demographic variation of 282 tree species at BCI, Panama (grey dots), is characterized by the growth–survival tradeoff (1st axis) distinguishing fast and slow species and a stature–recruitment tradeoff (2nd axis) distinguishing long-lived pioneers (LLP) and short-lived breeders (SLB). Arrows show loadings of a weighted PCA on annual diameter growth and survival rates of individuals ≥ 1 cm diameter in four canopy layers and the number of sapling recruits per unit of basal area. Colored dots are locations in demographic space of plant functional types that were used in model scenarios 1 and 3.

Our goal here is to explore whether this low-dimensional demographic tradeoff space can explain tropical forest dynamics, and if so, how much diversity along the tradeoff axes is necessary to accurately predict successional changes in species composition and basal area (a proxy for biomass carbon storage). We used species’ positions in the tradeoff space to back-calculate model parameters for all 282 species (*12*), thus smoothing across observed relationships between demographic rates. We simulated forest dynamics under four scenarios that differed in the number of tradeoffs (1 versus 2) and level of demographic diversity (number of simulated species or plant functional types (PFTs), Table 1, Fig. 2A).

**Table 1:** Model scenarios. Model scenarios differ in the number of included tradeoffs and level of demographic diversity (few PFTs vs 282 species). LLP – long-lived pioneers, SLB – short-lived breeders.

| Scenario | Tradeoffs | Demographic diversity |
| --- | --- | --- |
| 1 | Growth–Survival | 3 PFTs (fast, intermediate, slow) |
| 2 | Growth–Survival | 282 species |
| 3 | Growth–Survival, Stature–Recruitment | 5 PFTs (fast, slow, LLP, SLB, intermediate) |
| 4 | Growth–Survival, Stature–Recruitment | 282 species |

**Fig. 2:**
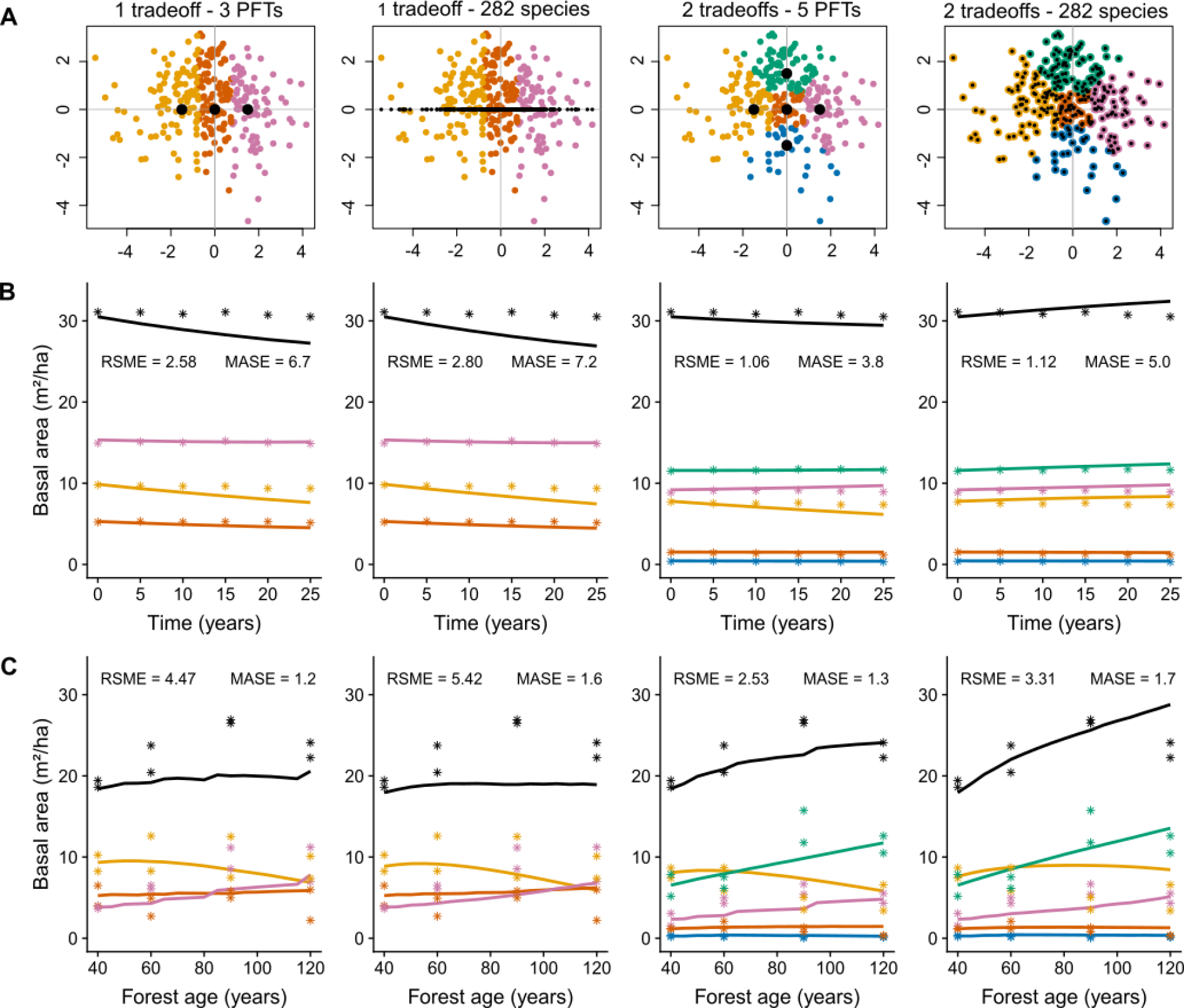
Predicted and observed basal area in four model scenarios (Table 1). (A) Locations of species (colored dots) and representative plant functional types (PFTs) or species used for model scenarios (black dots) in demographic space; each species was assigned to a PFT based on proximity in demographic space and color-coded as in Fig. 1. (B) Predicted (lines) and observed (asterisks) basal area by PFT in old-growth tropical forest (BCI, black is total basal area) and (C) secondary tropical forest (BCNM). RSME is the root mean square error of prediction of total basal area, MASE is the mean absolute scaled error of PFT-level predictions (*12*).

As a first test of predictive ability, we compared the observed dynamics of the 50-ha old-growth plot in BCI over 25 years with model predictions. We initialized the model with inventory data from 1985 and simulated forest dynamics until 2010. When only the growth– survival tradeoff was used, basal area was predicted to decline because of a decline of the number of trees >20 cm diameter, especially of fast species (Figs. 2B, S1). Including the stature– recruitment tradeoff axis improved the match between predicted and observed basal area (Fig. 2B) and above-ground biomass (AGB, Fig. S2) for different PFTs and size classes (Figs. S3, S4). However, when all species were simulated individually (scenario 4), the number of large trees (>60 cm diameter) and basal area were predicted to increase (Fig. S1). Maximum diameters were accurately predicted by all scenarios, except for scenario 2, where observed maximum diameters >150 cm were not reproduced (Fig. S5). The species-level parameterizations (scenarios 2 and 4) also reproduced the rank-abundance curve (Fig. S6).

As a second test of model performance, we initialized the model with data from 40-year old secondary forests in the Barro Colorado Nature Monument (BCNM) that are regenerating after abandonment from agricultural land use (*16*). We compared model predictions with observations from a chronosequence of 60, 90, and 120-year old secondary forests (two 1-ha plots in each age class). This is a strong test of model performance, because the model was parameterized with demographic data only from old-growth forest.

As in old-growth forest, predictions of secondary succession were most accurate when forest diversity was represented by 5 PFTs spanning both demographic tradeoffs. When only the growth–survival tradeoff was used, the increase of basal area (Fig. 2C) and AGB (Fig. S2) during succession was underestimated, because the number of large trees (>60 cm diameter) was underestimated (Fig. S7). In contrast, when both tradeoffs were included, observed successional changes in basal area, AGB, and abundance for different PFTs and size classes were accurately reproduced (Figs. 2C, S2, S7–S9). However, when all species were simulated individually (scenario 4), the number of large trees (>60 cm diameter) and basal area of fast species and LLPs was overestimated. The observed peak in basal area in the 90-year old secondary forest is likely caused by remnant trees in the study plots and disappears when larger spatial scales are considered (*17*). The diameter distribution after 400 years of simulation closely matched the observed diameter distribution only when both demographic tradeoffs were used (Fig. 3A).

**Fig. 3:**
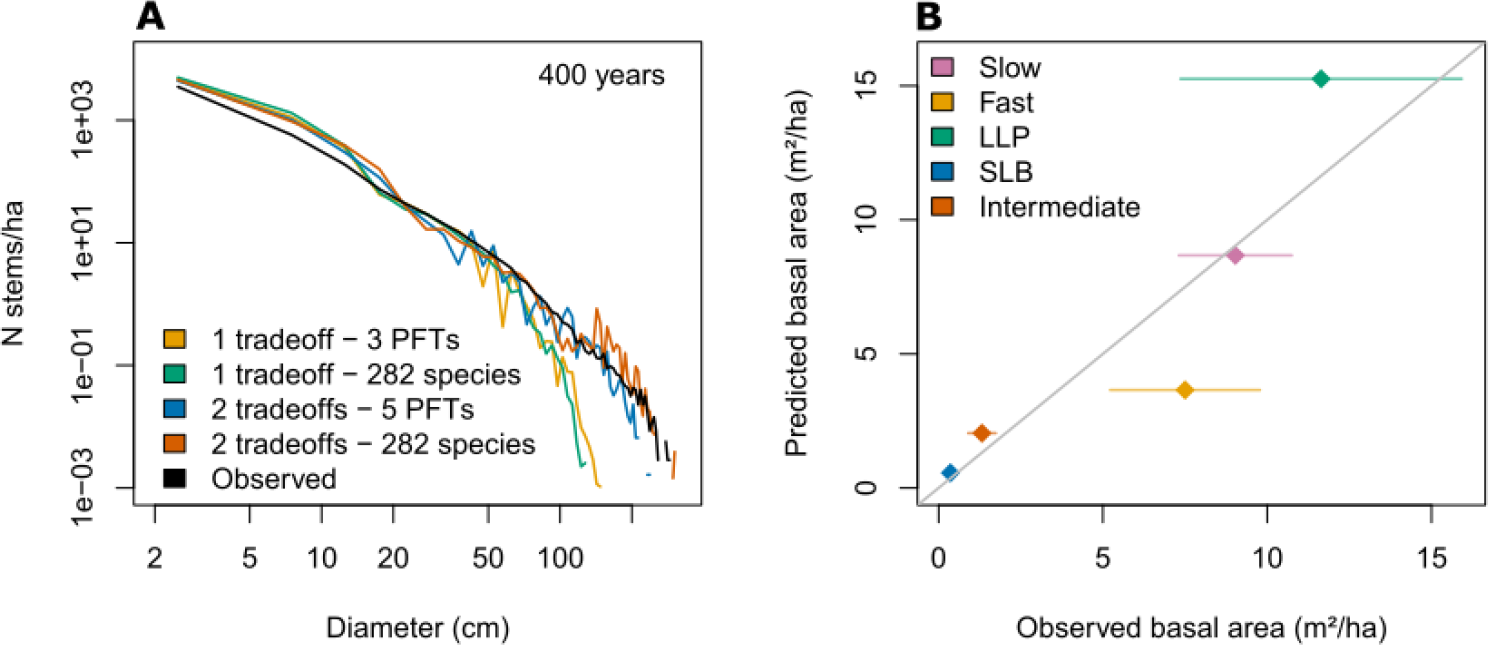
Model validation. (A) Diameter distribution in 400-year old simulated forest for the four model scenarios. (B) Predicted and observed basal area per PFT in model scenario 3 (5 PFTs spanning two demographic tradeoffs). Observed basal area is from an old-growth tropical forest in BCI, Panama. Predicted basal area is based on the estimated number of 0.1-ha patches in each age class (Fig. S11, *12*).

In a simulated long-term succession of scenario 3, slow species reached similar levels of basal area and AGB as LLPs after 400–500 years, and slow species and LLPs co-dominated the forest (Fig. S10). Fast species died out because the canopy gaps that they require for persistence (*18*) are treated in our model in a simplistic (non-spatially-explicit) manner. In reality, the forest is comprised of a mosaic of patches of different successional age since the last disturbance event (*19*). Comparing simulated successional trajectories of the fast and slow PFTs with observed species composition at the 0.1-ha scale allows inference of the patch-scale age distribution and suggests that the majority of the 0.1-ha patches within the BCI 50-ha plot are between 50 and 250 years old (Fig. S11, *12*). This model-inferred age distribution is consistent with LiDAR data collected on BCI, which suggest that between 0.43 and 1.6% of the area is disturbed every year, corresponding to an average disturbance interval between 63 and 233 years (*12, 20*). When we use the estimated proportion of 0.1-ha patches in each age class to generate the PFT-composition at equilibrium with the disturbance regime, predictions closely match observations (Fig. 3B).

Our results clearly show that two demographic tradeoffs are needed to accurately predict successional patterns in tropical forest structure and composition. Considering only the fast–slow continuum of life-histories is not sufficient, because it ignores long-lived pioneers, one of the most important (in terms of tree size and AGB) components in many tropical forests. Although the existence of long-lived pioneers has long been recognized (*5*), they have often been assumed to be part of the fast–slow continuum, i.e. considered to be mid-successional, because they reach their highest basal area in intermediate stages of succession (*6*). However, recent analyses show that long-lived pioneers lie on a second demographic dimension (*9*), and we demonstrate here for the first time that this second dimension is needed to accurately predict tropical forest dynamics. Understanding general patterns of tropical forest dynamics will require moving away from viewing succession through a one-dimensional to a two-dimensional lens of demographic strategies.

Our results suggest that the forest at BCI is in equilibrium with the local disturbance regime. This helps to resolve a long-standing dispute of whether long-lived pioneers are a transient feature of successional forests (*6, 21, 22*) and shows that, in this forest, they are not transient, but an integral and dominant component of the old-growth forest. In fact, long-lived pioneers dominate most successional stages and contribute more AGB than any other demographic group, except in very young forests (<40 years) or patches that have remained undisturbed for a long time (>400 years). Long-lived pioneers are able to maintain populations in the absence of large-scale disturbances and compensate for their low recruitment by growing quickly up to the canopy or emergent layer where they may remain as seed sources for several centuries (*9*).

Our results also suggest that a small number of demographic niches is sufficient to capture the dynamics of the BCI forest. Specifically, just 5 PFTs were sufficient to adequately capture successional patterns of forest composition and carbon dynamics. This result does not rule out the importance of additional functional axes (e.g., drought tolerance) at broader spatial or temporal scales, or the potential importance of individual species for ecosystem functioning. Nevertheless, our results suggest that functional diversity in species-rich tropical forests may be much smaller than taxonomic diversity, and that tropical forest diversity could be accurately represented in Earth System Models by a small number of PFTs that span the relevant functional axes (*23*).

This systematic assessment of the optimal demographic resolution was only possible because all model parameters are directly observable in the field, and because tradeoffs in model parameters capture the relevant dimensions of functional variation among species, with no need for perfect information for each species (*23, 24*). Most importantly, the close match between field observations and the definition of model parameters avoids the necessity for time-consuming and error-prone manual parameter tuning (*25*) or computationally-intensive inverse parameter fitting (*26*). More complex tropical forest models require data that are not available for most species, e.g. light transmission, light-use efficiency, respiration rates, seed-dispersal kernels, germination rates etc. (*27*). While such details can provide mechanistic insights, our simplified approach, based on observed recruitment rates, and growth, and mortality rates in different canopy layers, allowed for direct tests of hypotheses about demographic tradeoffs and functional diversity (e.g. 5 PFTs vs many species) in structuring tropical forest dynamics.

Together, the demographic forest model and the empirical demographic tradeoffs define an objective, reproducible, and automated workflow that scales up from demographic rates of individual species to community structure and dynamics. Given its ease of application and the increasing availability of tropical forest inventory data, this workflow has the potential to substantially advance theoretical understanding of tropical forest dynamics by comparing demographic diversity, structure, and dynamics of forests that differ in climate, floristic composition, and/or disturbance regime. It also has the potential to facilitate the evidence-based planning of forest restoration and sustainable tropical forest management by providing improved quantitative tools for predicting rates and trajectories of forest regrowth (*4, 9*). Lastly, the two-dimensional tradeoff studied here could also be combined with other key axes (e.g. drought tolerance) in a low-dimensional tradeoff space, allowing for improved representation of tropical forest functional diversity in Earth System Models (*23*).

## Materials and Methods

### The PPA model

We used a version of the PPA model that is based on Purves et al. (*11*), where tree crowns are assumed to be flat. The simulation area was 1 ha and the model time step was 5 years. The model works on cohorts of trees that share the same age, diameter at breast height (dbh, in cm) and species/plant functional type (PFT). The number of trees in a cohort can be fractions of individuals, including numbers <1. Cohorts are removed from the simulation when they have <0.001 individuals. We extended the model from two to four canopy layers (*14*) and species/PFTs are characterized by growth and mortality rates in each of the four layers. We modified several aspects of the model. Cohorts are removed if they are assigned to a layer >4. Sapling cohorts enter the model at 1 cm dbh (originally 0.01 cm). Recruitment rates are constant (see below, originally they scaled with a species’ crown area in the canopy layer). Sapling cohorts recruit to layer 4. The dbh (cm)-crown radius (m) relationship is nonlinear (originally linear),

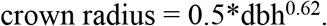

Likewise, the dbh (cm)-height (m) relationship is non-linear and parameters for both allometries were determined using data from BCI (*28*),

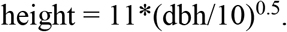

As a single allometry for all trees worked equally well as species-specific allometries in determining structural and dynamics properties of the forest (*14*), we used a single allometry for crown radius and height.

To calculate above-ground biomass (AGB, Mg), we followed ForestGEO protocols and used allometric equations based only on dbh and wood density (wd), but not height, from Chave et al. (*29*) for moist tropical forest:

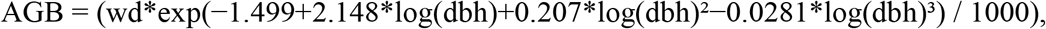

where dbh is measured in cm and wd in g/cm^3^.

### Parameterization

In a previous study, we performed a weighted PCA (*15*) on nine demographic parameters for 282 species from the BCI 50-ha plot, namely growth rate in the four canopy layers, survival (expressed as lifespan) in the four canopy layers, and the number of recruits per unit of adult basal area, which were derived from forest inventory data (*9, 30*). We follow the taxonomy as of 2017 (*31*). The first two principal components of this PCA correspond to the fast–slow continuum (37% explained variation) and a stature–recruitment tradeoff (28% explained variation), respectively. Here we used a slightly modified version of the PCA using the number of recruits per unit of total species’ basal area, and used the first two principal components (henceforth ‘axes’ or ‘tradeoffs’) to determine model parameters. An exception are recruitment rates, which we determined directly from forest inventory data (independent of the basal area of a species and independent from the PCA). We assumed recruitment rates to be constant over time because the 50-ha plot is embedded within a larger forest area from which seeds continuously arrive into the study area. Moreover, relationships between recruitment rates per PFT and total basal area in 31.25 × 31.25 m^2^ subplots or basal area of the respective PFT were weak or absent (not shown).

To determine growth and mortality rates, we specified coordinates of five PFTs symmetrically in the two-dimensional demographic space (Fig. 1 in main manuscript):

- Intermediate (location x_1_=0, x_2_=0)
- Fast (location x_1_=−1.5, x_2_=0)
- Slow (location x_1_=1.5, x_2_=0)
- Long-lived pioneer (LLP, location x_1_=0, x_2_=1.5)
- Short-lived breeder (SLB, location x_1_=0, x_2_=−1.5)

Coordinates of +/−1.5 on the two tradeoff axes correspond to between 9 and 19% of species having more extreme demographic strategies.

For the simulations including all species, we used their PCA scores along the 1^st^ or 1^st^ and 2^nd^ PCA tradeoff axis, depending on the scenario.

We then solved the linear system of equations consisting of the PCA loadings of the nine parameters (Table S1) and species’ scores (setting all species’ scores on axes 3 to 9 to 0, i.e. x_3_…x_9_ = 0) to obtain transformed input parameters to the PCA. These were then back-transformed to model parameters by de-centering, de-scaling, and de-logging. Lifespan was transformed into mortality, i.e. mortality = 1/lifespan (Tables S2,S3).

From these strategies, we simulated four scenarios, differing in the number of species/PFTs:

1. 1 tradeoff, 3 PFTs (fast, intermediate, slow)
2. 1 tradeoff, 282 species
3. 2 tradeoffs, 5 PFTs (slow, fast, LLP, SLB, intermediate)
4. 2 tradeoffs, 282 species.

Annual recruitment rates (at 1 cm dbh) for each PFT were determined as the average number of recruits (per ha) of species that were assigned to the PFT. For scenarios (2) and (4), species without observed recruits (25 species) were assigned one recruit in 25 years and 50 ha, i.e. 0.0008 recruits per year and ha. New recruits enter the simulation every year and experience deterministic mortality every year. However, annual recruit numbers were determined from 5-year census intervals. Thus, we adjusted annual recruit numbers by species/PFT-specific mortality such that, after a 5-year time step, simulated recruit numbers matched observed average recruit numbers in 5-year census intervals in the 50-ha plot at BCI.

Wood density (wd) for PFTs was determined as the volume-weighted mean of wd in old-growth forest. Wood density is from bci.spptable (*32*; sometimes to genus or family level only). Individual tree volume was calculated as

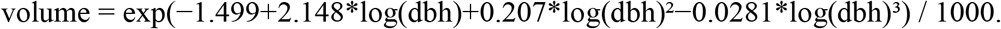

Volume-weighted wood density of the PFTs in secondary forest was slightly different from that of the PFTs in the old-growth forest, due to differences in species’ abundance. We used the volume-weighted wood density of the PFTs in old-growth forest, when we calculated AGB in simulations of old-growth forest dynamics, and wood density of the PFTs in secondary forest plots, when we calculated AGB in simulations of secondary forest succession.

### Species assignment to PFTs, model initialization and validation

#### Old-growth forest

In the 50-ha permanent plot in tropical moist forest on Barro Colorado Island (BCI), Panama, every tree ≥ 1 cm dbh is tagged, mapped, and measured approximately every five years (*30*). In this paper our analyses are based on six censuses (conducted between 1985 and 2010). We leave out the first census of 1982 because in this census some tall trees with buttresses were measured at lower heights than in subsequent censuses introducing a bias in basal area and AGB estimates. Detailed methods for the plot censuses can be found in (*2*) and (*13*).

For comparison of model predictions with data, we assigned species to PFTs based on their PCA scores along the 1^st^ or 1^st^ and 2^nd^ PCA axis. For scenarios (1) and (3), we assigned species to the PFT with the closest location that was used for parameterization (Figs. 1, 2A in main manuscript). For scenario (1), 98 species were assigned to the ‘fast’ PFT, 83 to the ‘slow’ PFT, and 101 to the ‘intermediate’ PFT. For scenario (3), 75 species were assigned to the ‘fast’ PFT, 76 to the ‘LLP’ PFT, 60 to the ‘slow’ PFT, 30 to the ‘SLB’ PFT, and 41 to the ‘intermediate’ PFT. For visualization purposes, we used the same PFT assignments for scenarios (2) and (4), where all species were simulated individually.

For simulation of old-growth forest dynamics, we initialized the model with the average (in terms of species abundances and tree sizes) of the 50-ha plot on Barro Colorado Island in 1985. Individuals of species that were not included in the PCA (mostly palms and hemiepiphytes, 1.4% of individuals, 3.5% of basal area) were omitted in these calculations as they could not be associated with a PFT. Thus, the initial state of the model is slightly less populated than the real forest. Species were assigned to one of 111 size classes and tree numbers were aggregated by size class and species/PFT. Size classes were 1 cm wide for individuals between 1 and 50 cm dbh, 2 cm wide for individuals between 50 and 100 cm dbh, and 5 cm wide for larger individuals. The lower limit of the size class was used as initial cohort size in the PPA model.

We validated the model against field data in terms of overall basal area, AGB, and abundance per PFT, as well as in different size classes. Forest structure and composition was determined from the six censuses of the 50-ha plot (1985–2010). Basal area and AGB were compared for the size classes 1–20 cm, 20–60 cm, ≥ 60 cm, ≥ 1 cm dbh (total). Abundance was compared for the size classes 5–20 cm, 20–60 cm, ≥ 60 cm, ≥ 5 cm dbh (total).

As measures of predictive power, we calculated the root mean square error (RMSE) of prediction for total basal area. RMSE measures the average deviation of the predicted value from the observed value and is in the same unit as observations (m^2^/ha). We also calculated the mean absolute scaled error (MASE) to compare the predictive power of different model parameterizations at the PFT level that are at different scales (*33*). MASE is scale-independent and measures the predictive power of a model relative to a naïve random walk forecast.

We compared simulated (after 100 years of simulation) and observed maximum diameters. Maximum diameter for each PFT in the field data and the simulations was calculated as the largest 5-cm diameter class with >0.1 individuals per ha. For parameterizations (2) and (4) (282 species), it was calculated for each species as the largest 5-cm diameter class with >0.005 individuals per ha. For parameterizations (2) and (4) (282 species), we additionally compared simulated and observed rank-abundance curves. Observed abundance was calculated as average abundance per ha over six censuses (1985–2010). We used the ‘RADanalysis’ package in R (*34*) to construct a rank abundance distribution using the max rank normalization method.

We compared the diameter distribution of simulated 400-year old forest with the observed diameter distribution of the old-growth forest. The observed diameter distribution again is an average of six censuses and includes palms and hemiepiphytes.

#### Secondary forest

Data on secondary forests is from eight forest plots (1 ha each) <7 km away from the old-growth forest plot (*16, 35, 36*). There were two plots in each of four age classes (40, 60, 90, and 120 years). All secondary forest stands had been in agriculture, including pasture, swidden farming, and plantation farming, for undetermined lengths of time prior to fallow (*16*). The plots were inventoried between 2011 and 2014. In all plots, every tree ≥ 5 cm dbh was tagged, mapped, and measured, and in most of the plots, in a 0.5-ha subset of the plot every tree ≥ 1 cm dbh was tagged, mapped, and measured. We only considered the largest stem of multi-stemmed individuals to match old-growth forest data.

We excluded 254 individuals without recorded dbh as well as palms (8 species, 474 individuals), hemiepiphytes (1 species, 4 individuals), cultivated species (1 species, 1 individual), and unidentified individuals (214). Of the remaining 242 species (9935 individuals), we had no information on demographic strategy from the old-growth forest for 50 species (1212 individuals). We assigned some of these species to the PFTs of a closely related species, and others based on average demographic characteristics of taxonomically-related species and/or species with similar functional traits, i.e. wood density and growth form. Wood density and growth form is from bci.spptable (*32*; sometimes to genus or family level only).

For simulation of secondary forest succession, we initiated the model with the average of two 1-ha 40-year old secondary forest plots. Species were assigned to one of 111 size classes and tree numbers were aggregated by size class and PFT. Size classes were 1 cm wide for individuals between 1 and 50 cm dbh, 2 cm wide for individuals between 50 and 100 cm dbh, and 5 cm wide for larger individuals. The lower limit of the size class was used as initial cohort size in the PPA model.

We validated the model against field data in terms of total basal area, AGB, and abundance per PFT, as well as in different size classes. Basal area, AGB, and abundance were compared for the size classes 5–20 cm, 20–60 cm, ≥ 60 cm, ≥ 5 cm dbh (total), because sampling of the different secondary forest plots <5 cm dbh was inconsistent. To calculate RSME and MASE, we averaged the observations in the two 1-ha plots per age class to yield a single time series of basal area.

### Disturbance regime

Analyses from detailed LiDAR data from the year 2009 estimated 0.43% of the area of BCI to be canopy gaps with <2 m canopy height and 1.6% of the area to be canopy gaps <5 m canopy height (*20*). Assuming that the vegetation can re-grow to a canopy height between 2 and 5 m within one year, the fraction of the forest that is disturbed every year is between 0.43 and 1.6%. This corresponds to an average disturbance interval between 62.5 (100/1.6) and 232.6 (100/0.43) years.

### Age distribution and simulated equilibrium mixed forest

We divided inventory data from the six censuses between 1985 and 2010 from the 50-ha plot into 512 31.25 m × 31.25 m subplots and calculated the basal area (m^2^/ha) of species assigned to the slow and fast PFTs of scenario (3) for each subplot. Then, we determined the year (in steps of 5 years) in a simulated succession to which the basal area of fast and slow PFTs in each subplot was most similar, respectively, and took their mean. As the model was initialized with inventory data from 40-year old forest, we linearly extrapolated the basal area of PFTs for younger ages between 0 m^2^/ha for year 0 and the observed basal area at year 40. The resulting combined (across censuses) age distribution of subplots (Fig. S11) was then used to generate a ‘simulated’ equilibrium of the mixed forest as the sum of simulated basal area or AGB of the respective ages, weighted by the proportion of subplots in the respective age class. Note: We only considered the fast and slow PFTs because they show a clear successional pattern, while LLPs maintain high and SLBs and the intermediate PFT maintain low basal area throughout much of the succession.

## Acknowledgments

We thank D. Purves for sharing the PPA model code and D. Purves, S. A. Bohlman and H. C. Muller-Landau for inspiring discussions and helpful suggestions;

## Author contributions

N.R. designed the research with input from C.F., J.L., and C.W. N.R. and C.F. performed the research. R.C., D.D., S.D., S.H., and O.L. provided data. N.R. drafted the first version of the paper, and all authors contributed to subsequent versions of the paper.

## Supplementary Figures and Tables

**Fig. S1.**
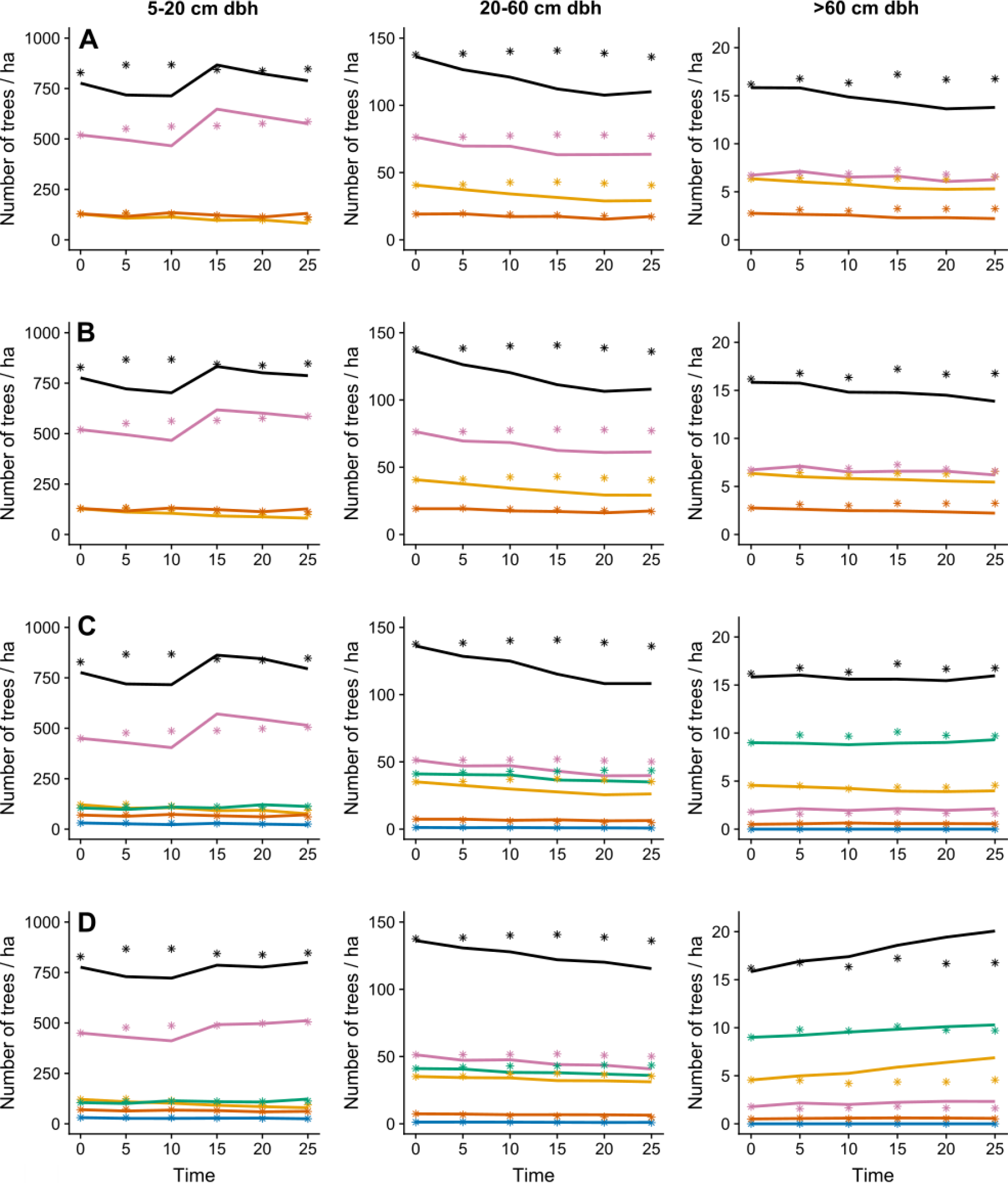
Predicted and observed abundance in four model scenarios (rows; A: 1 tradeoff – 3 PFTs, B: 1 tradeoff – 282 species, C: 2 tradeoffs – 5 PFTs, D: 2 tradeoffs – 282 species) and three size classes (columns). Simulated (lines) and observed (asterisks) abundance per PFT in an old-growth tropical forest in Barro Colorado Island, Panama. Color code: purple – slow, yellow – fast, green – LLP, blue – SLB, red – intermediate, black – total.

**Fig. S2.**
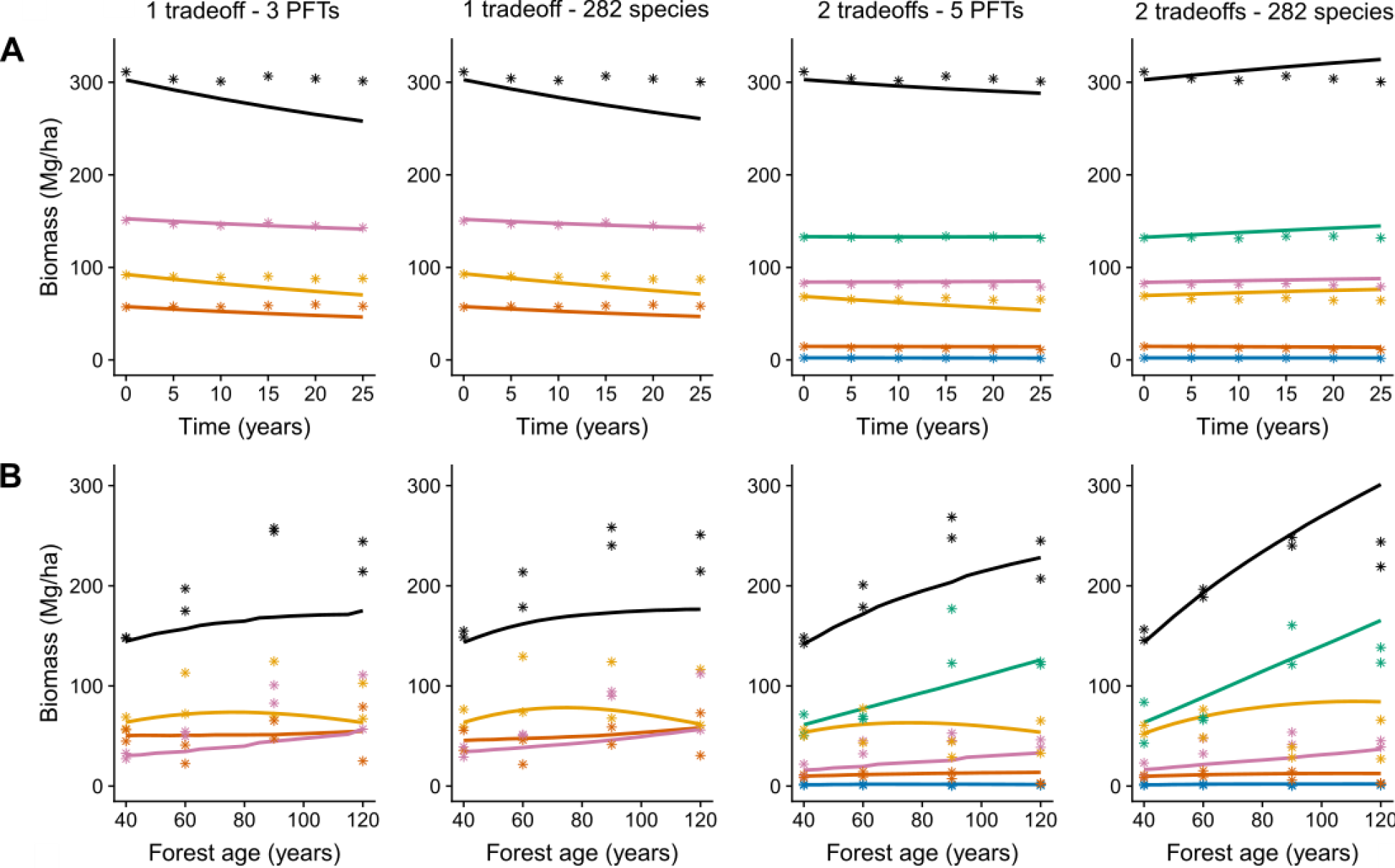
Predicted and observed aboveground biomass (AGB) in four model scenarios that differ in the number and demographic characteristics of simulated species or PFTs. (A) Predicted (lines) and observed (asterisks) biomass per PFT in an old-growth tropical forest in Barro Colorado Island, Panama (≥ 1 cm dbh) and (B) in secondary tropical forest in Barro Colorado Nature Monument National Park, Panama (≥ 5 cm dbh). Color code: purple – slow, yellow – fast, green – LLP, blue – SLB, red – intermediate, black – total.

**Fig. S3.**
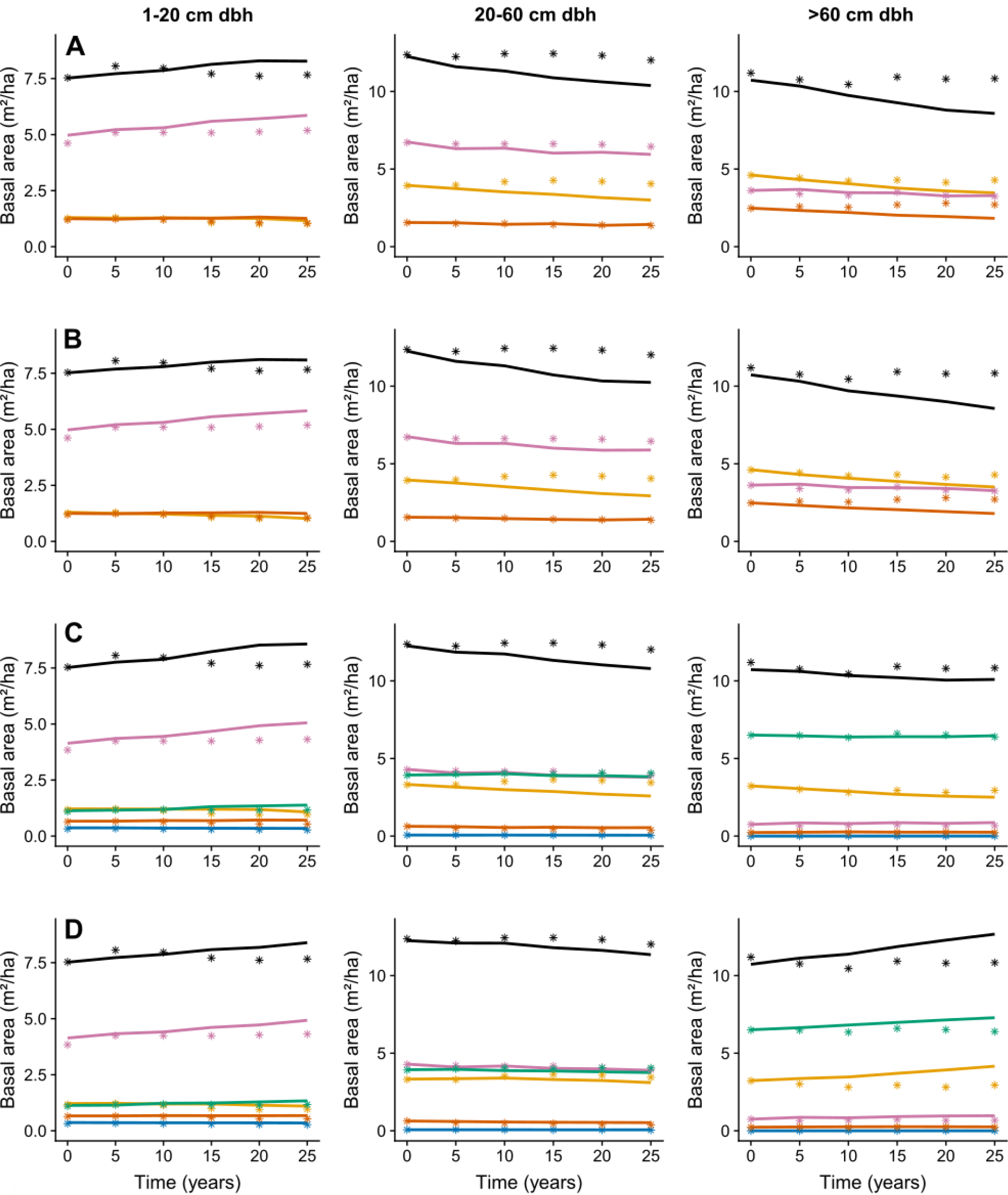
Predicted and observed basal area in four model scenarios (rows; A: 1 tradeoff – 3 PFTs, B: 1 tradeoff – 282 species, C: 2 tradeoffs – 5 PFTs, D: 2 tradeoffs – 282 species) and three size classes (columns). Simulated (lines) and observed (asterisks) basal area per PFT in an old-growth tropical forest in Barro Colorado Island. Color code: purple – slow, yellow – fast, green – LLP, blue – SLB, red – intermediate, black – total.

**Fig. S4.**
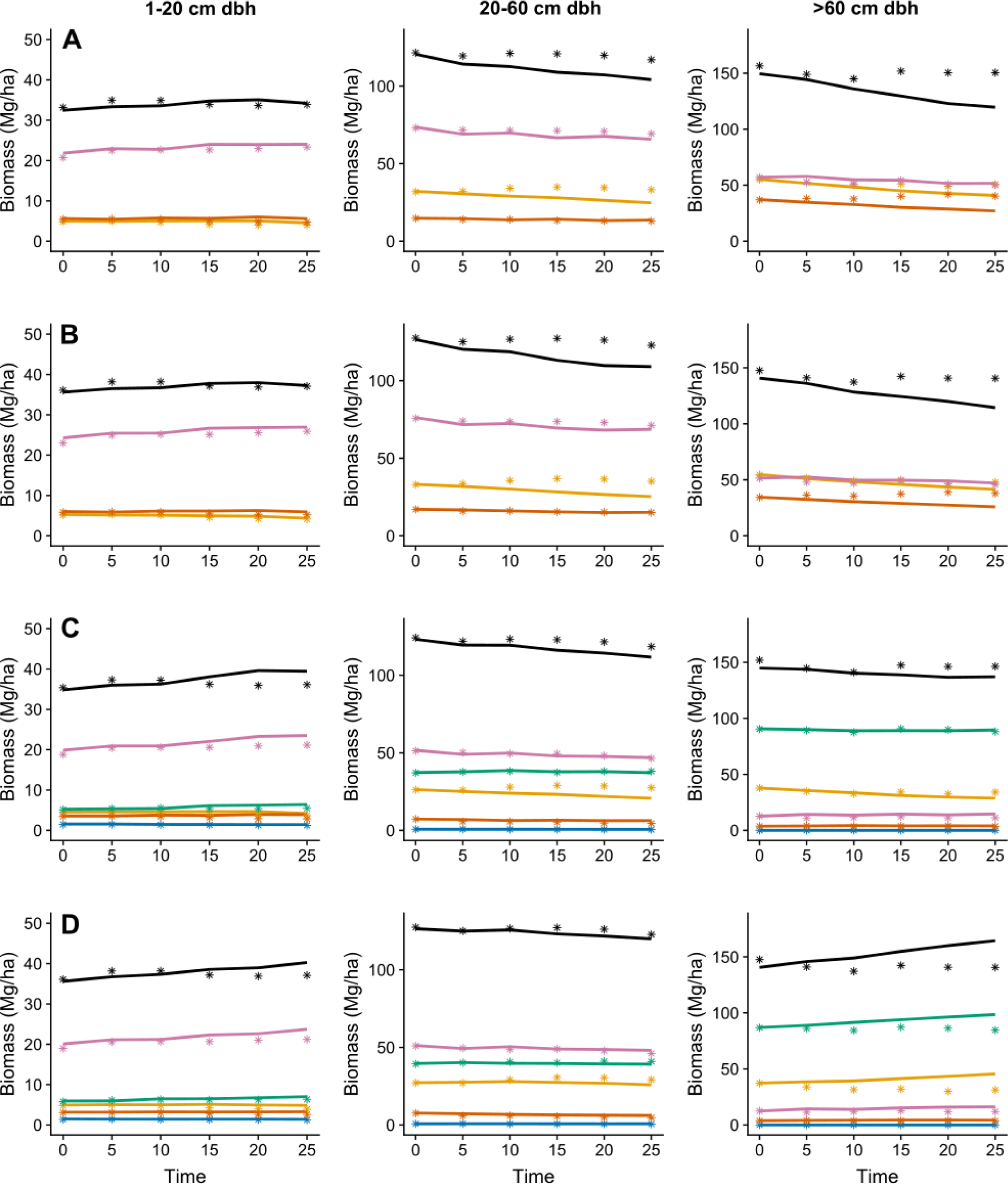
Predicted and observed above-ground biomass (AGB) in four model scenarios (rows; A: 1 tradeoff – 3 PFTs, B: 1 tradeoff – 282 species, C: 2 tradeoffs – 5 PFTs, D: 2 tradeoffs – 282 species) and three size classes (columns). Simulated (lines) and observed (asterisks) biomass per PFT in an old-growth tropical forest in Barro Colorado Island, Panama. Color code: purple – slow, yellow – fast, green – LLP, blue – SLB, red – intermediate, black – total.

**Fig. S5.**
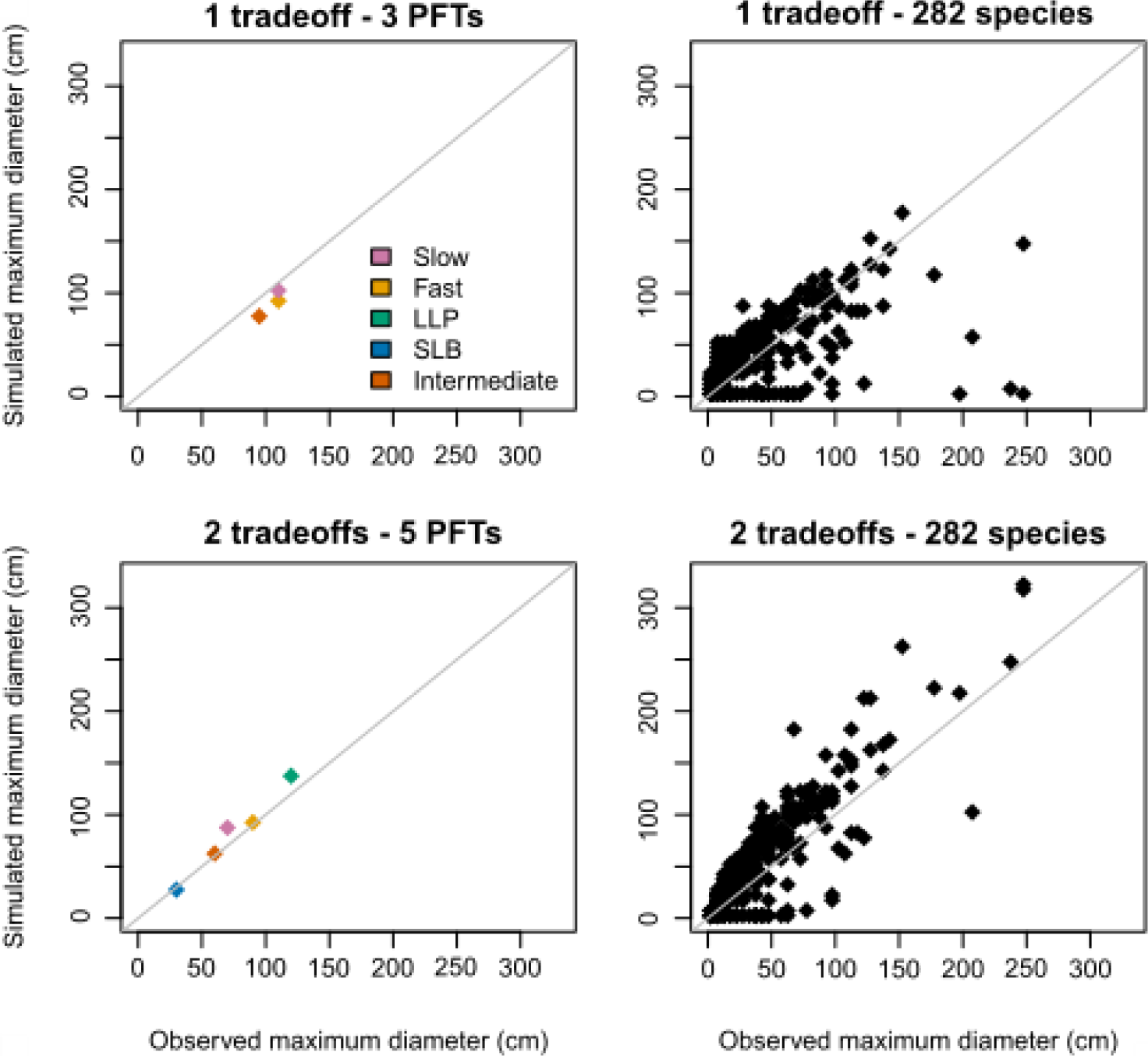
Simulated and observed maximum diameters of the PFTs or species for four model scenarios. 1:1 lines are shown in grey. See Suppl. for details.

**Fig. S6.**
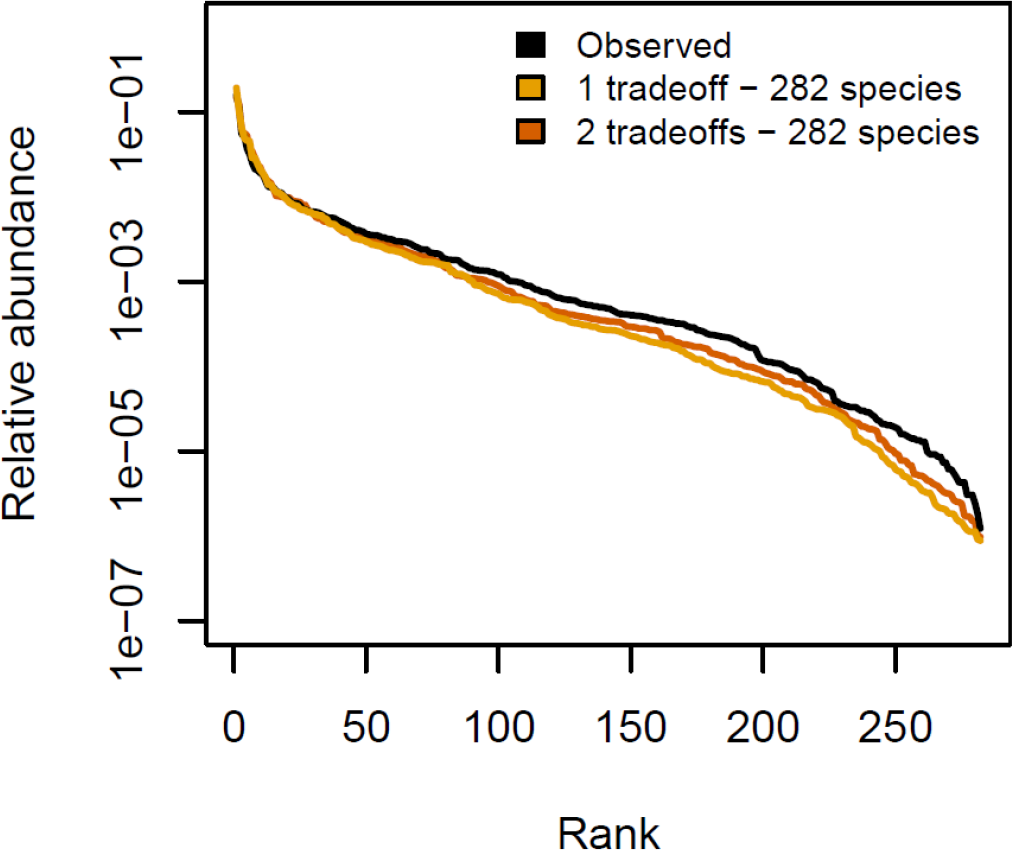
Observed and predicted rank-abundance curves after 100 years of simulation of old-growth forest at Barro Colorado Island, Panama.

**Fig. S7.**
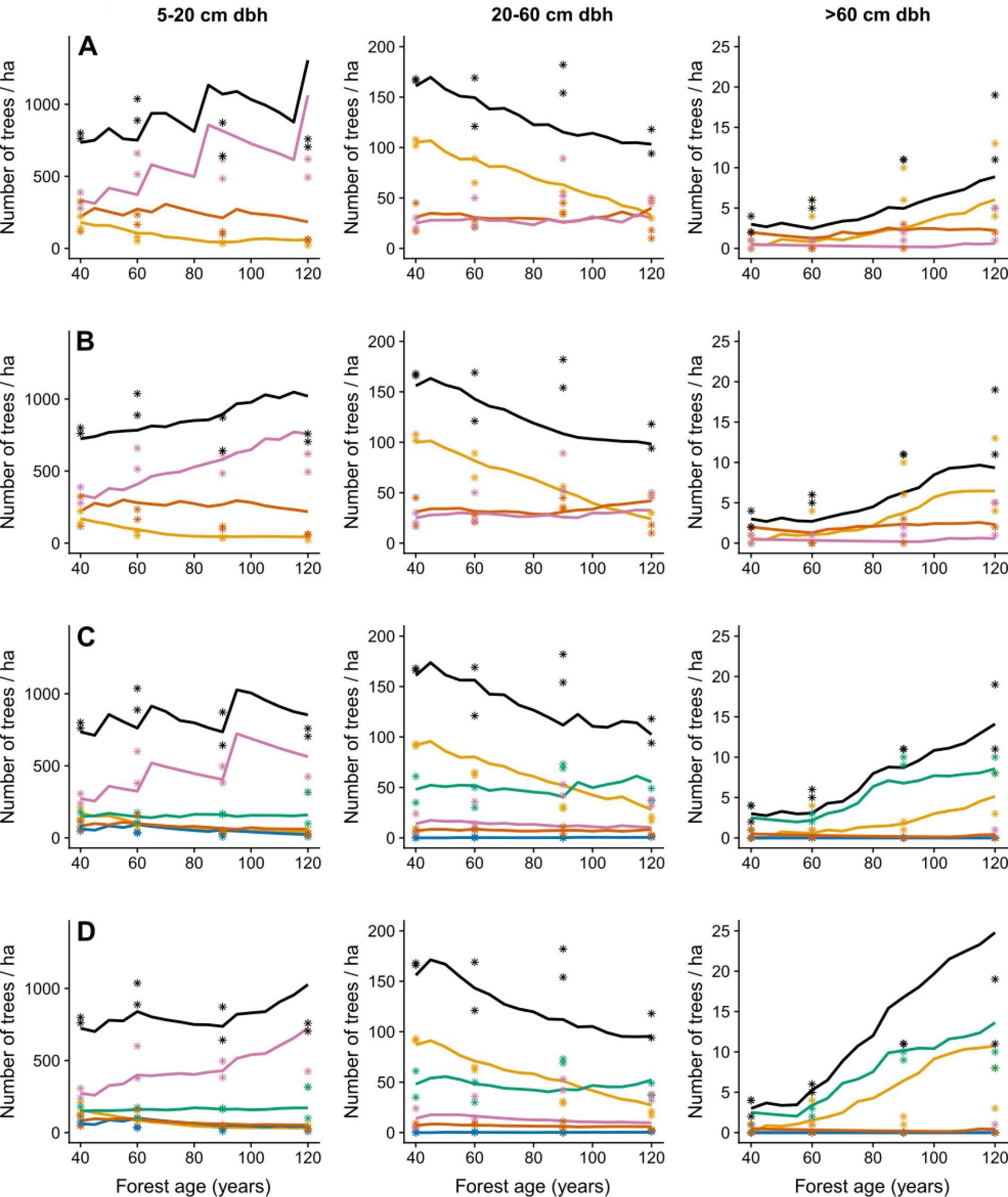
Predicted and observed abundance in four model scenarios (rows; A: 1 tradeoff – 3 PFTs, B: 1 tradeoff – 282 species, C: 2 tradeoffs – 5 PFTs, D: 2 tradeoffs – 282 species) and three size classes (columns). Simulated (lines) and observed (asterisks) abundance per PFT in secondary tropical forest in Barro Colorado Nature Monument National Park, Panama. Color code: purple – slow, yellow – fast, green – LLP, blue – SLB, red – intermediate, black – total.

**Fig. S8.**
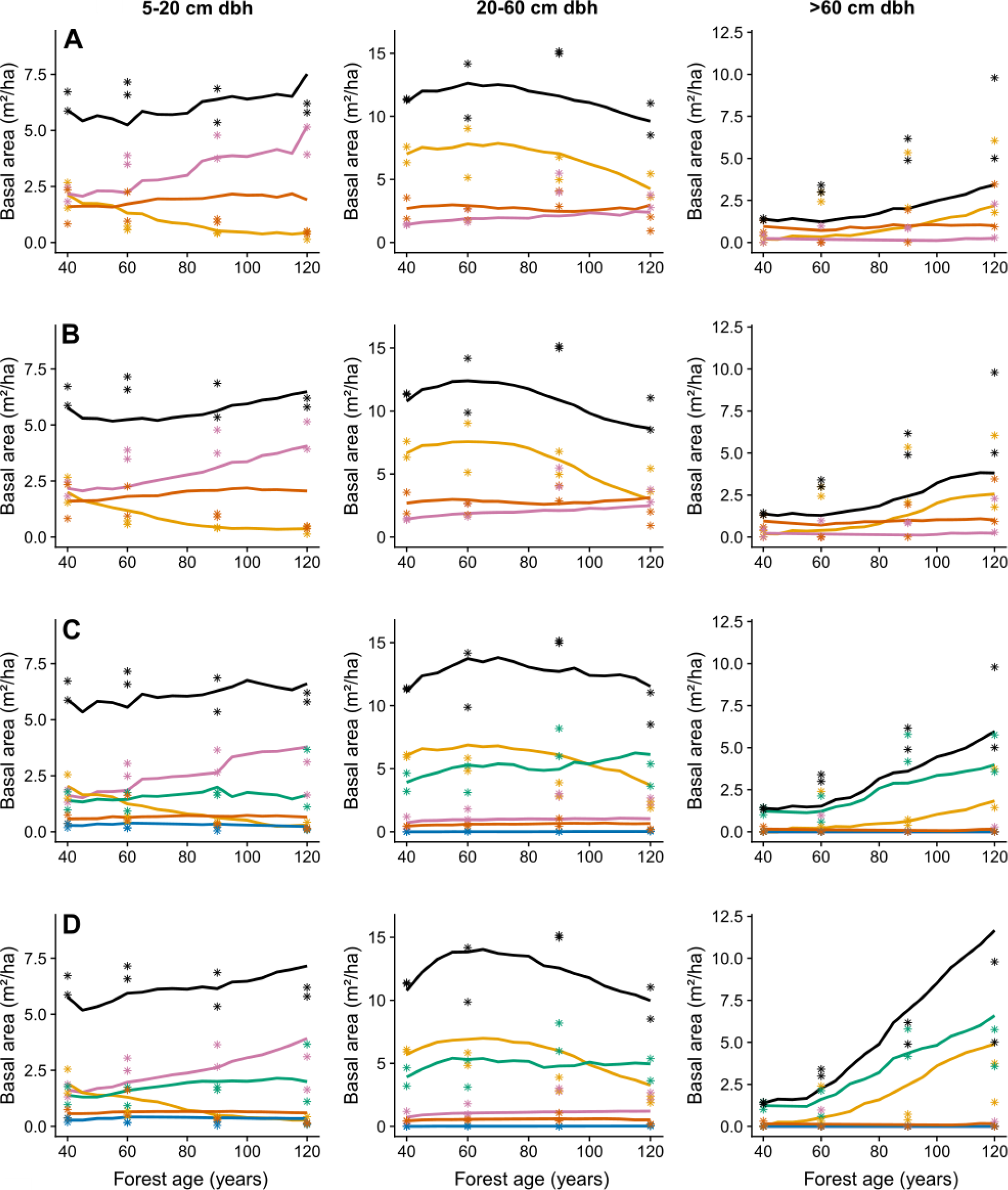
Predicted and observed basal area in four model scenarios (rows; A: 1 tradeoff – 3 PFTs, B: 1 tradeoff – 282 species, C: 2 tradeoffs – 5 PFTs, D: 2 tradeoffs – 282 species) and three size classes (columns). Simulated (lines) and observed (asterisks) basal area per PFT in secondary tropical forest in Barro Colorado Nature Monument National Park, Panama. Color code: purple – slow, yellow – fast, green – LLP, blue – SLB, red – intermediate, black – total.

**Fig. S9.**
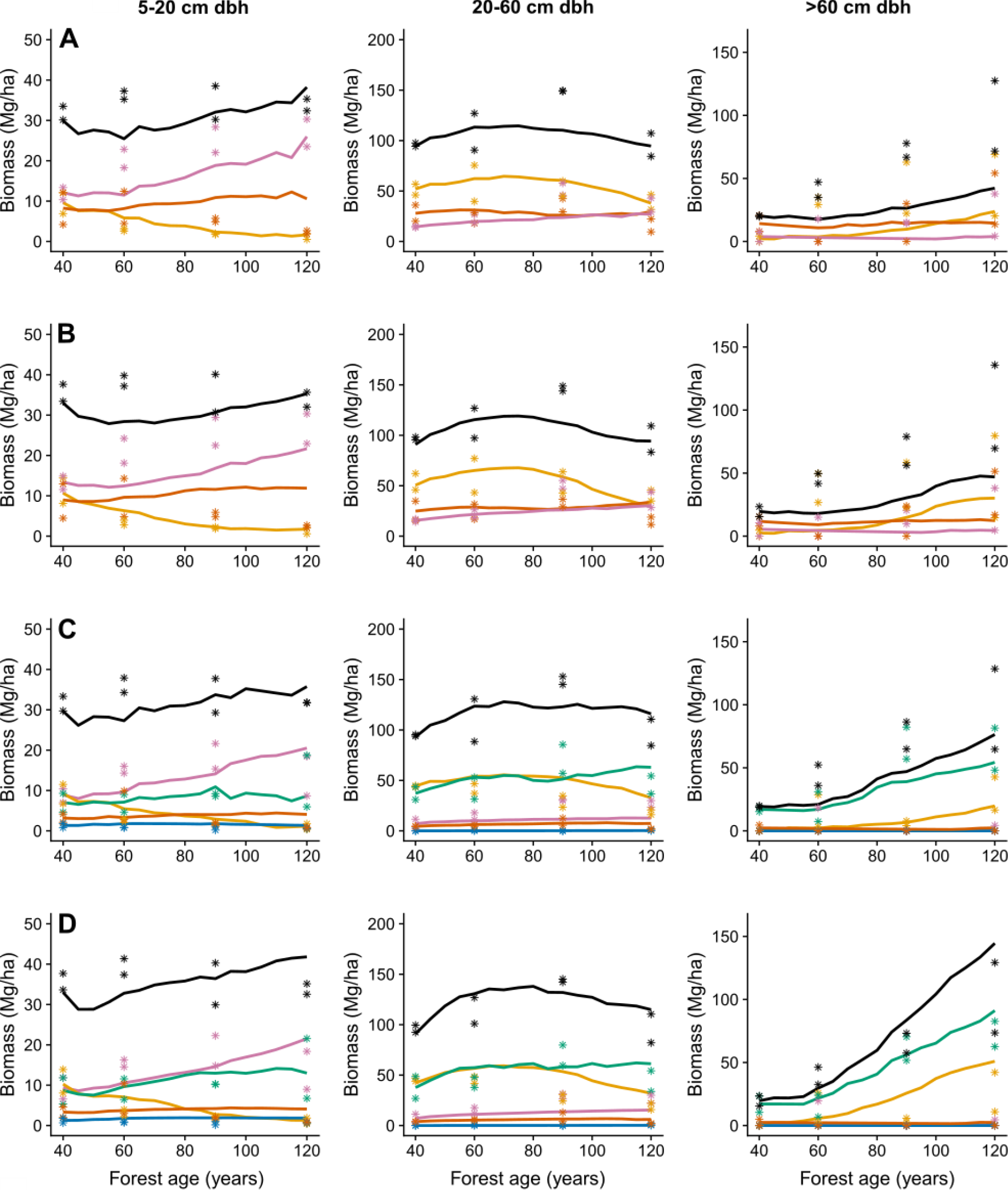
Predicted and observed above-ground biomass (AGB) area in four model scenarios (rows; A: 1 tradeoff – 3 PFTs, B: 1 tradeoff – 282 species, C: 2 tradeoffs – 5 PFTs, D: 2 tradeoffs – 282 species) and three size classes (columns). Simulated (lines) and observed (asterisks) biomass per PFT in secondary tropical forest in Barro Colorado Nature Monument National Park, Panama. Color code: purple – slow, yellow – fast, green – LLP, blue – SLB, red – intermediate, black – total.

**Fig. S10.**
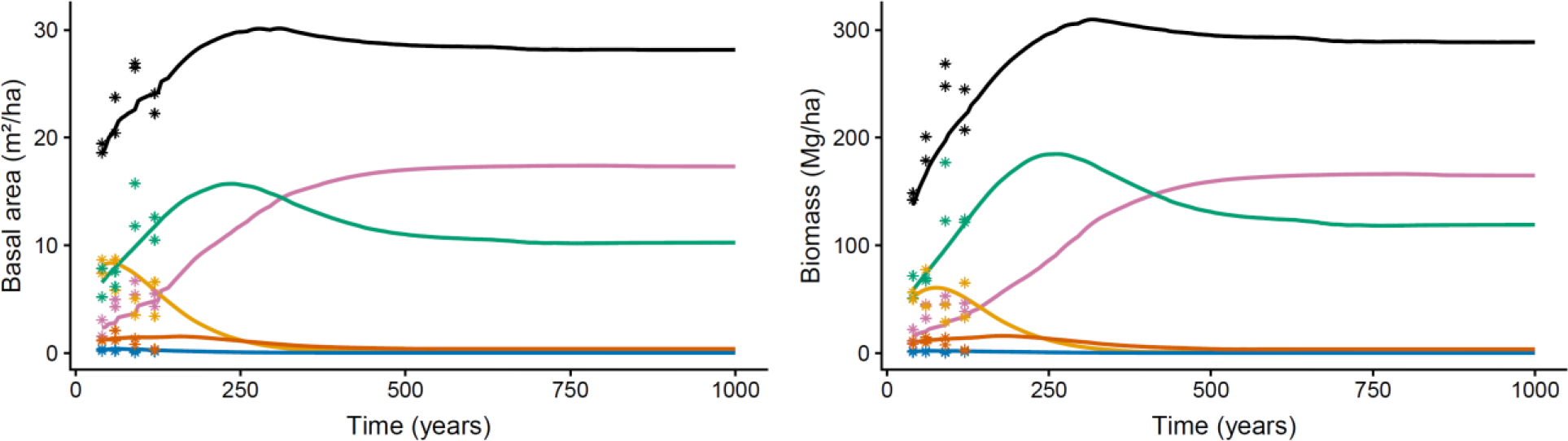
Simulated long-term succession of scenario 3 (2 tradeoffs, 5 PFTS) starting from inventory data of 40-year old secondary forest. Basal area and above-ground biomass of individuals ≥ 5 cm dbh for comparison with secondary forest data (asterisks). Recruitment rates of the PFTs are constant and set to annual averages of the number of observed recruits of species assigned to the five PFTs. Color code: purple – slow, yellow – fast, green – LLP, blue – SLB, red – intermediate, black – total.

**Fig. S11.**
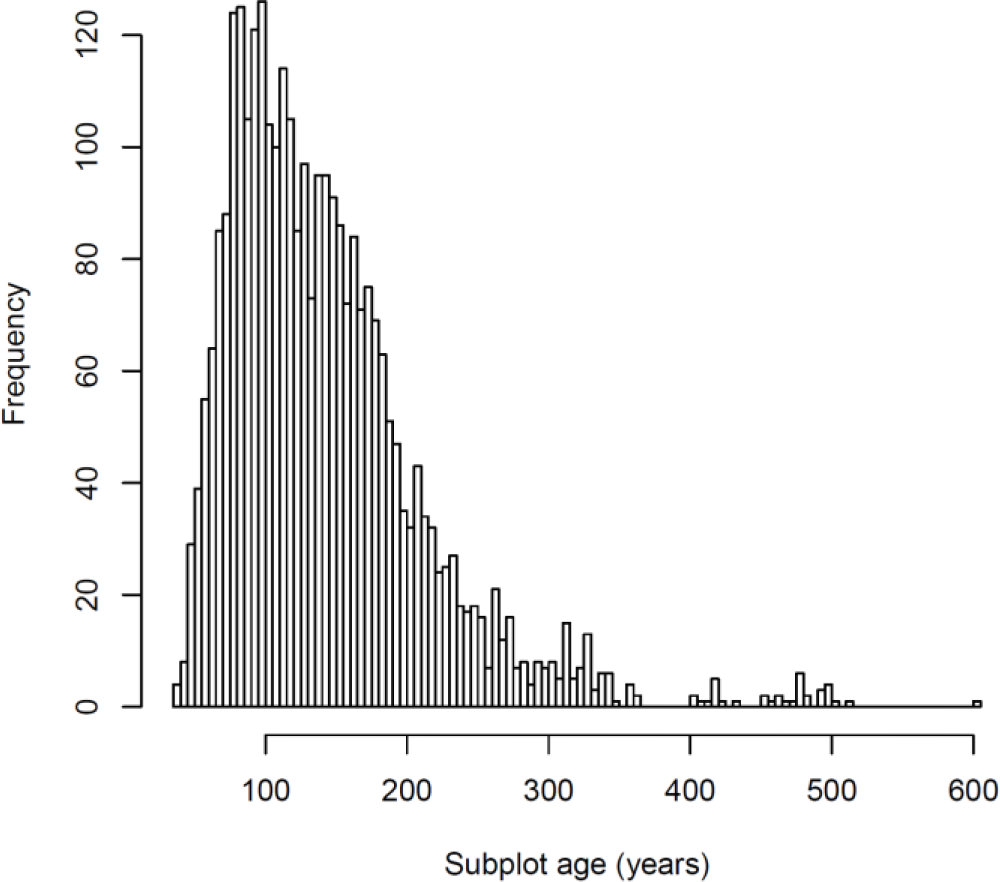
Estimated age distribution of ~0.1-ha subplots from six censuses (1985, 1990, 1995, 2000, 2005, 2010).

**Table S1.** Loadings of demographic parameters in the wPCA. Survival and growth of trees (≥ 1 cm dbh) in four canopy layers is indicated by ‘Survival 1’ etc. Recruitment is the number of recruits per unit of total species basal area. Only the first one or two principal components are used to back-calculate model parameters from species scores in PCA space, depending on the scenario.

| Parameters | Parameter loadings |  |  |  |  |  |  |  |  |
| --- | --- | --- | --- | --- | --- | --- | --- | --- | --- |
|  | PC1 | PC2 | PC3 | PC4 | PC5 | PC6 | PC7 | PC8 | PC9 |
| Survival1 | 0.2103 | 0.2947 | -0.5398 | -0.4130 | -0.4612 | 0.3415 | -0.2380 | 0.1407 | -0.0356 |
| Survival2 | 0.3370 | 0.4051 | -0.0063 | -0.2672 | 0.3237 | 0.0814 | 0.7127 | -0.0482 | 0.1712 |
| Survival3 | 0.4376 | 0.2396 | 0.2830 | 0.2505 | 0.0200 | 0.1136 | -0.4280 | -0.1090 | 0.6323 |
| Survival4 | 0.4541 | 0.2032 | 0.2832 | 0.2428 | 0.1178 | 0.2067 | -0.1305 | 0.1782 | -0.7127 |
| Growth1 | -0.1943 | 0.2070 | -0.6090 | 0.5235 | 0.4575 | 0.2204 | -0.0746 | -0.1046 | -0.0014 |
| Growth2 | -0.3504 | 0.1586 | 0.2375 | -0.5400 | 0.5401 | 0.2434 | -0.3889 | 0.0320 | -0.0285 |
| Growth3 | -0.3899 | 0.3775 | 0.1968 | 0.2308 | -0.1839 | 0.0735 | 0.1434 | 0.7262 | 0.1622 |
| Growth4 | -0.3609 | 0.3588 | 0.2738 | 0.1188 | -0.3651 | 0.3276 | 0.1296 | -0.6163 | -0.1215 |
| Recruitment | 0.0415 | -0.5578 | 0.0858 | 0.0571 | 0.0069 | 0.7750 | 0.2025 | 0.1228 | 0.1404 |

**Table S2.**
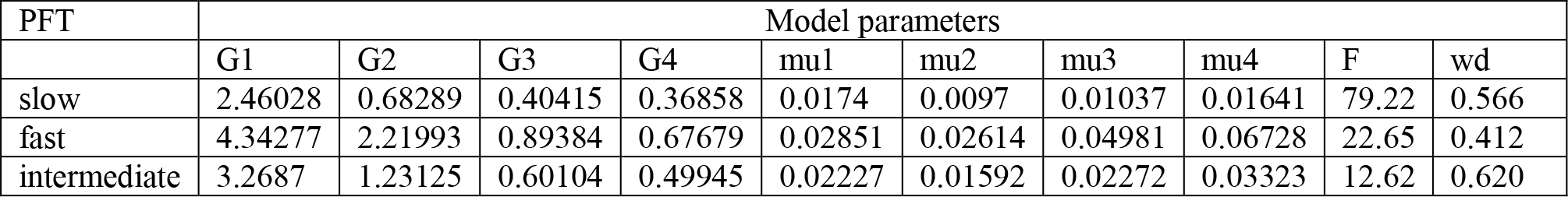
PPA model parameters for 3 PFTs (1 demographic tradeoff axis). G1 to G4 and mu1 to mu4 are annual growth (mm/y) and mortality (1/y) rates in four canopy layers, respectively. F is the number of new recruits over the 1 cm dbh threshold per year and hectare and wd is wood density (g/cm³).

**Table S3.** PPA model parameters for 5 PFTs (2 demographic tradeoff axes). G1 to G4 and mu1 to mu4 are annual growth (mm/y) and mortality (1/y) rates in four canopy layers, respectively. F is the number of new recruits over the 1 cm dbh threshold per year and hectare and wd is wood density (g/cm³).

| PFT | Model parameters |  |  |  |  |  |  |  |  |  |
| --- | --- | --- | --- | --- | --- | --- | --- | --- | --- | --- |
|  | G1 | G2 | G3 | G4 | mu1 | mu2 | mu3 | mu4 | F | wd |
| slow | 2.46028 | 0.68289 | 0.40415 | 0.36858 | 0.0174 | 0.0097 | 0.01037 | 0.01641 | 65.37 | 0.635 |
| Fast | 4.34277 | 2.21993 | 0.89384 | 0.67679 | 0.02851 | 0.02614 | 0.04981 | 0.06728 | 20.83 | 0.403 |
| LLP | 4.42383 | 1.6079 | 0.88258 | 0.67557 | 0.01576 | 0.00877 | 0.01479 | 0.02423 | 6.22 | 0.480 |
| SLB | 2.4152 | 0.94283 | 0.40931 | 0.36925 | 0.03148 | 0.0289 | 0.03492 | 0.04557 | 16.83 | 0.653 |
| intermediate | 3.2687 | 1.23125 | 0.60104 | 0.49945 | 0.02227 | 0.01592 | 0.02272 | 0.03323 | 6.24 | 0.600 |

